# Dysregulated synaptic gene expression in oligodendrocytes of spinal and bulbar muscular atrophy

**DOI:** 10.1101/2024.01.11.575248

**Authors:** Madoka Iida, Kentaro Sahashi, Tomoki Hirunagi, Kenji Sakakibara, Kentaro Maeda, Yosuke Ogura, Masaki Iizuka, Tomohiro Akashi, Kunihiko Hinohara, Masahisa Katsuno

## Abstract

Spinal and bulbar muscular atrophy (SBMA) is a neuromuscular disease caused by an expanded CAG repeat in the *androgen receptor* (*AR*) gene. To elucidate the cell type-specific temporal gene expression in SBMA, we performed single-nucleus RNA sequencing on the spinal cords of AR-97Q mice. Among all cell types, oligodendrocytes (OLs) had the highest number of differentially expressed genes before disease onset. Analysis of OL clusters suggested that pathways associated with cation channels and synaptic function were activated before disease onset, with increased output from OLs to neurons in AR-97Q mice compared to wild-type mice. These changes in the early stages were abrogated in the advanced stages. An OL cell model of SBMA showed phenotypes similar to those of AR-97Q mice at early stages, such as increased transcriptional changes in synapse organization. Our results indicate that the dysregulation of cell-to-cell communication has a major impact on the early pathology of SBMA and is a potential therapeutic target for SBMA.

## Introduction

Spinal and bulbar muscular atrophy (SBMA) is an X-linked, adult-onset neuromuscular disease caused by a CAG repeat expansion within the first exon of the *androgen receptor* (*AR*) gene (William R. Kennedy *et al*, 1968). It is characterized by progressive muscle weakness, atrophy, and fasciculation of the limb and bulbar muscles, which manifest between 30 and 60 years of age (Hashizume *et al*, 2020). Serum creatinine concentrations are significantly reduced in patients with SBMA and correlate with disease severity (Lombardi *et al*, 2019). MRI assessment of skeletal muscle and fat is another promising biomarker for tracking disease changes (Klickovic *et al*, 2019). Although the ligand-dependent toxicity of the polyglutamine-expanded AR protein is central to the pathogenesis of SBMA, the initiation and progression of its degenerative processes remain elusive.

As in other neurodegenerative disorders (McDade *et al*, 2021; Kilzheimer *et al*, 2019), preclinical changes have been identified in SBMA. Most patients with SBMA notice hand tremors and muscle cramps more than 10 years before the emergence of limb weakness (Hijikata *et al*, 2018). Patients present with elevated serum creatine kinase levels and changes in muscle pathology before or at the onset of clinical symptoms (Sorarù *et al*, 2008). Female carriers of SBMA may develop mild muscular weakness associated with changes in neurogenic biomarkers such as decreased motor unit number estimation and electromyographic abnormalities (Torii *et al*, 2023). Several mouse models of SBMA show reduced muscle force and altered contractile properties as early pathological events (Gray *et al*, 2020). It has also been demonstrated that the skeletal muscles of SBMA knock-in mice show metabolic changes such as increased lipid metabolism and impaired glycolysis prior to denervation (Rocchi *et al*, 2016). Furthermore, presymptomatic SBMA transgenic mice (AR-100Q) show early changes in the expression pattern of genes involved in muscle contraction and structure (Marchioretti *et al*, 2023).

In addition to the degeneration of motor neurons and skeletal muscle (Iida *et al*, 2015, 2019; Yu *et al*, 2006; Cortes *et al*, 2014; Sahashi *et al*, 2015; Hirunagi *et al*, 2023), glial cell alterations are also observed in certain mouse models of SBMA. The limb muscles of AR-113Q mice exhibit myopathy-like features and reduced mRNA levels of neurotrophin-4 and glial cell-derived neurotrophic factor, suggesting that glial cells are involved in the development of SBMA (Yu *et al*, 2006). TGF-β signaling, which plays a crucial role in the survival and function of adult neurons, was dysregulated in motor neurons as well as glial cells of SBMA model AR-97Q transgenic mice (Katsuno *et al*, 2010). It has also been reported that astrocyte proliferation is prominent and inflammatory M1 microglia are prevalent in the spinal cord of AR-97Q mice (Iida *et al*, 2015).

To understand the early pathogenesis of SBMA and to systematically assess the role of different cell types in the central nervous system of SBMA, we examined gene expression in the spinal cord of AR-97Q mice at the single-nucleus level during different stages of the disease, including before the onset of motor dysfunction. Oligodendrocytes (OLs) had the highest number of differentially expressed genes (DEGs) from the preonset phase, and the expression of the genes related to ion channel and synapse function in OLs was upregulated in the early stages but downregulated in the advanced stage of SBMA.

## Results

### Single-nucleus sequencing of the spinal cord from SBMA model AR-97Q mice revealed transcriptional alterations in oligodendrocytes (OLs)

To uncover cell type-specific temporal transcriptional changes, we conducted single-nucleus RNA sequencing (snRNA-seq) on spinal cord samples from AR-97Q mice and wild-type mice at four stages: prepubertal (3 weeks of age), preonset (6 weeks of age), early symptomatic (9 weeks of age) and advanced (13 weeks of age) (n = 4 mice for each condition) (Fig. 1A). Transcriptional profiles of an average of 8800 nuclei per sample were obtained (Supplementary Fig. 1A, B). The average total transcript reads across conditions was over 20,000 reads per cell. After applying quality filters, 54,456 cells were retained for further analysis (Supplementary Fig. 1C). Samples from all timepoints were combined and clustered together to obtain an overview of the complete spinal cord cell dataset. Data were projected onto two dimensions via t-distributed stochastic neighbor embedding (t-SNE) and uniform manifold approximation and projection (UMAP) (Fig. 1B). Cell type designations were first determined by analyzing the DEGs in each cluster and manually comparing them to several canonical markers of each cell type (Supplementary Fig. 2A). The major cell classes in the sequencing dataset showed clear segregation according to the previously reported markers of these cell types, and 10 major cell classes were identified: oligodendrocytes (OLs), inhibitory neurons, excitatory neurons, astrocytes, oligodendrocyte precursor cells (OPCs), OL progenitors, microglia, pericytes, endothelial cells, and fibroblasts. The OL progenitors, known as committed oligodendrocyte progenitors (COPs), differed from OPCs in that they lacked *Pdgfra* and *Cspg4* and expressed *Neu4* and genes expressed by undifferentiated oligodendrocytes, such as *Sox6*, *Bmp4*, and *Gpr17* (Marques *et al*, 2016) (Supplementary Fig. 2B). The results revealed that the proportion of OLs was the highest compared to that of each cell type within the samples, consistent with a previous report from human and mouse spinal cords (Yadav *et al*, 2023; Zeisel *et al*, 2018) (Fig. 1C). The proportion of OLs was lower in AR-97Q mice than in wild-type mice, while the proportion of neurons was higher in AR-97Q mice than in wild-type mice at all ages. The number of DEGs in the OLs of AR-97Q mice compared to wild-type mice at 3, 6, and 9 weeks was the highest number of DEGs among all cell types (Fig. 1D–F). At 13 weeks, OLs had the second highest number of DEGs after microglia (Fig. 1G). Color-coding of the snRNA-seq data by weeks of age also showed that the gene expression of OLs in AR-97Q mice began to change from 3 weeks of age, with the differences from wild-type mice becoming more pronounced as time progressed (Fig. 1H).

**Fig. 1.**
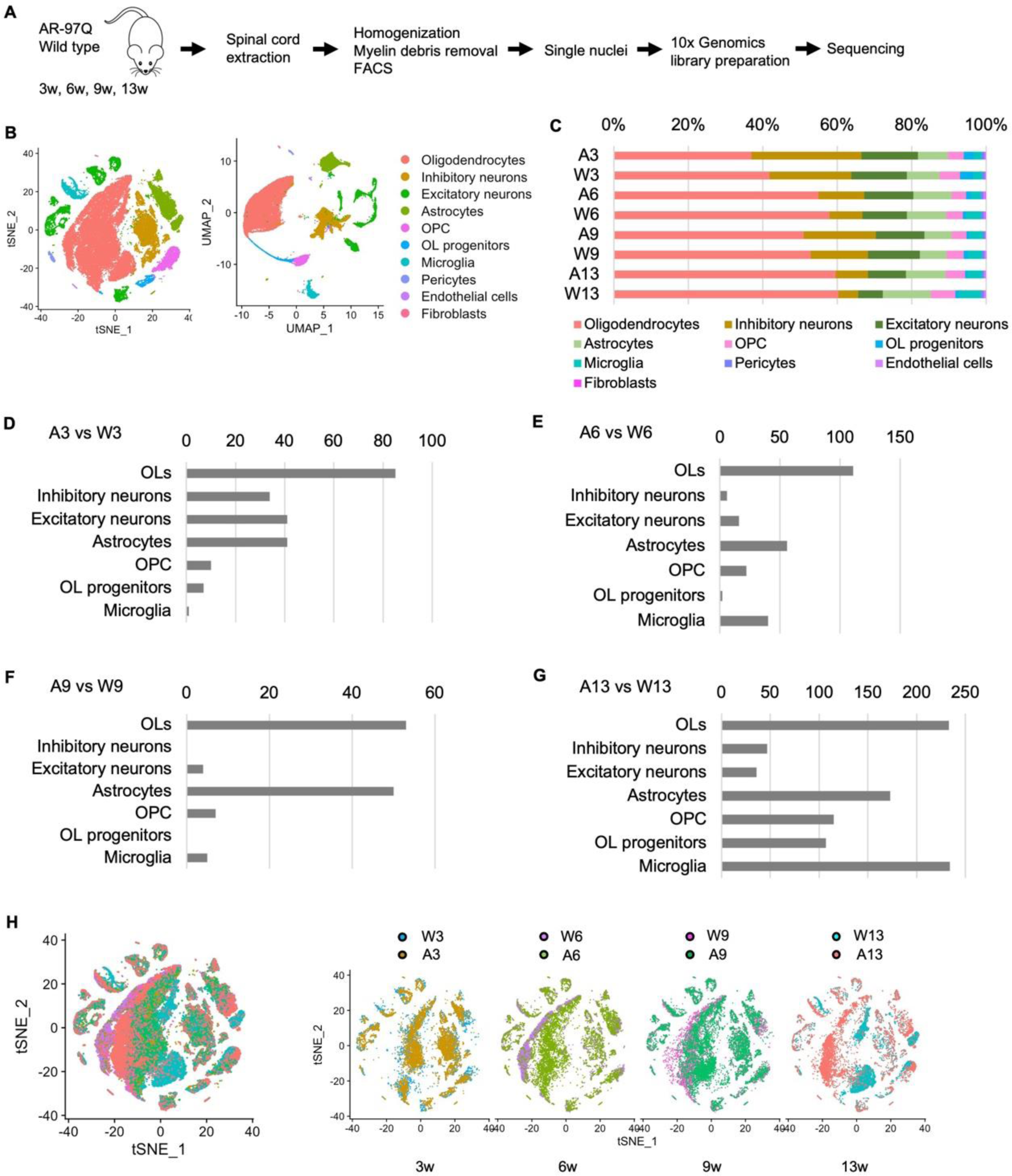
Cell type-specific differences between AR-97Q and wild-type mice according to single-nucleus RNA sequencing (snRNA-seq) **A,** Experimental workflow for snRNA-seq analysis. **B,** t-Distributed stochastic neighbor embedding (t-SNE) and uniform manifold approximation and projection (UMAP) plots of 54,456 nuclei from the spinal cords of all mice used in this experiment. **C,** Proportion of the 9 cell types of each sample. **D–G,** Number of DEGs in each cell type in AR-97Q mice and wild-type mice. Adjusted *p* value < 0.05, |log_2_FC| ≥ 0.20 for 3 weeks of age (**D**) and |log_2_FC| ≥ 0.40 for 6, 9, and 13 weeks of age (**E–G**). **H**, t-SNE plot of nuclei color-coded by each sample (left) and t-SNE plots of nuclei comparing AR-97Q mice with wild-type mice at four different disease stages (right). A, AR-97Q mice; W, wild-type mice (ex. A3 indicates AR-97Q mice at 3 weeks); OLs, oligodendrocytes. N = 4 mice for each sample.

Human AR was expressed in glial cells in the spinal cord of AR-97Q mice at 3 weeks, and immunofluorescence staining indicated that OLs also express human AR (Fig. 2A, B). Immunohistochemical staining with the 1C2 antibody, which specifically recognizes expanded polyglutamine, showed 1C2-positive cells in the spinal cords of AR-97Q mice at 6, 9, and 13 weeks, whereas no 1C2-positive cells were observed at 3 weeks (Fig. 2C). Immunoblotting analyses revealed that the expression level of Sox10, a marker of OPCs and OLs, was significantly lower in the spinal cord of AR-97Q mice than in that of wild-type mice at 13 weeks (Fig. 2D, E). The immunoreactivity of myelin basic protein (Mbp), a marker of myelin, was lower in AR-97Q mice than in wild-type mice at 13 weeks (Supplementary Fig. 3). We further examined the implication of OLs in human SBMA pathology. Polyglutamine inclusions were present in OLs in autopsy specimens of the spinal cord from patients with SBMA (Fig. 2F). Immunoblotting analyses showed that the level of SOX10 was significantly lower in the spinal cord of SBMA patients than in control individuals (Fig. 2G, H). MBP in the autopsy spinal cord specimens of SBMA patients had lower immunoreactivity than MBP in control patients (Fig. 2I), suggesting that OLs are impaired in the spinal cords of patients with SBMA.

**Fig. 2.**
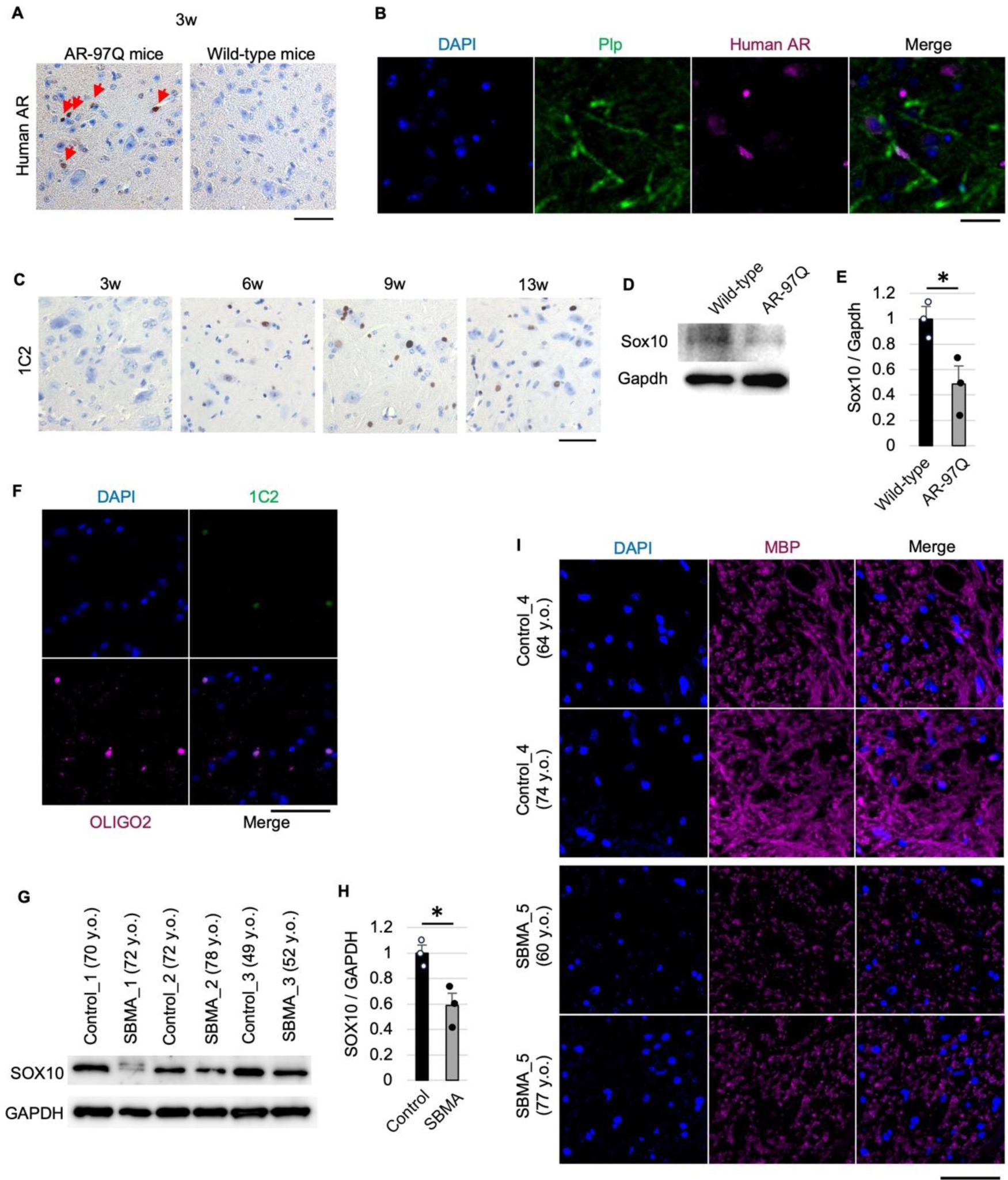
Impaired oligodendrocytes (OLs) in AR-97Q mice and patients with SBMA. **A,** Expression of human androgen receptor (*AR*) in the glial cells of AR-97Q mice at 3 weeks. The arrows indicate glial cells expressing human AR. **B,** Expression of human AR in the OLs of AR-97Q mice at 3 weeks. **C,** Immunohistochemical staining of the spinal cord from 3-, 6-, 9-, and 13-week-old AR-97Q mice with an antibody against polyglutamine (1C2). **D,** Immunoblotting of Sox10 in the spinal cords of wild-type and AR-97Q mice. **E,** Quantitative analysis of the expression level of Sox10 in the spinal cords of wild-type and AR-97Q mice (n = 3 mice per group). **F,** Expression of polyglutamine in the OLs of the spinal cord from patients with SBMA. **G,** Immunoblotting of SOX10 in autopsied spinal cords in disease controls and SBMA subjects. All subjects were males. **H,** Quantitative analysis of the expression level of SOX10 in the spinal cords of control and SBMA subjects (n = 3 subjects per group). **I,** Immunohistochemical analysis of MBP in the autopsied spinal cords of control and SBMA subjects. Error bars indicate the SEM. \**p* < 0.05, unpaired two-sided t test. Scale bar: 50 μm (**A**). Scale bars: 50 μm (**A, F, I**) or 25 μm (**B, C**). y.o., years old.

### Differentially expressed genes at each week

To investigate the transcriptional changes in OLs before the appearance of motor symptoms, we compared the data obtained from AR-97Q and wild-type mice at 6 weeks and specified 10 major cell types by unsupervised clustering (Fig. 3A). The volcano plot of DEGs from the OL cluster of AR-97Q and wild-type mice showed that the difference between the two groups was evident (Fig. 3B, C). The predicted protein interaction (PPI) networks for the top 20 DEGs upregulated in AR-97Q mice showed that they are related to each other and involve several genes associated with the cation channel complex and synaptic membrane, although such interaction was unclear for downregulated genes (Fig. 3D, E). The Gene Ontology (GO) analysis of the top 100 upregulated DEGs in AR-97Q mice (log_2_FC > 0.404) revealed that the DEGs were associated with ion channel activity and synapse organization in the GO biological process and molecular function categories (Fig. 3F, G). In contrast, the top 100 downregulated genes in AR-97Q mice (log_2_FC < −0.224) were associated with tubulin binding, actin binding, and positive regulation of cell projection organization (Fig. 3H, I).

**Fig. 3.**
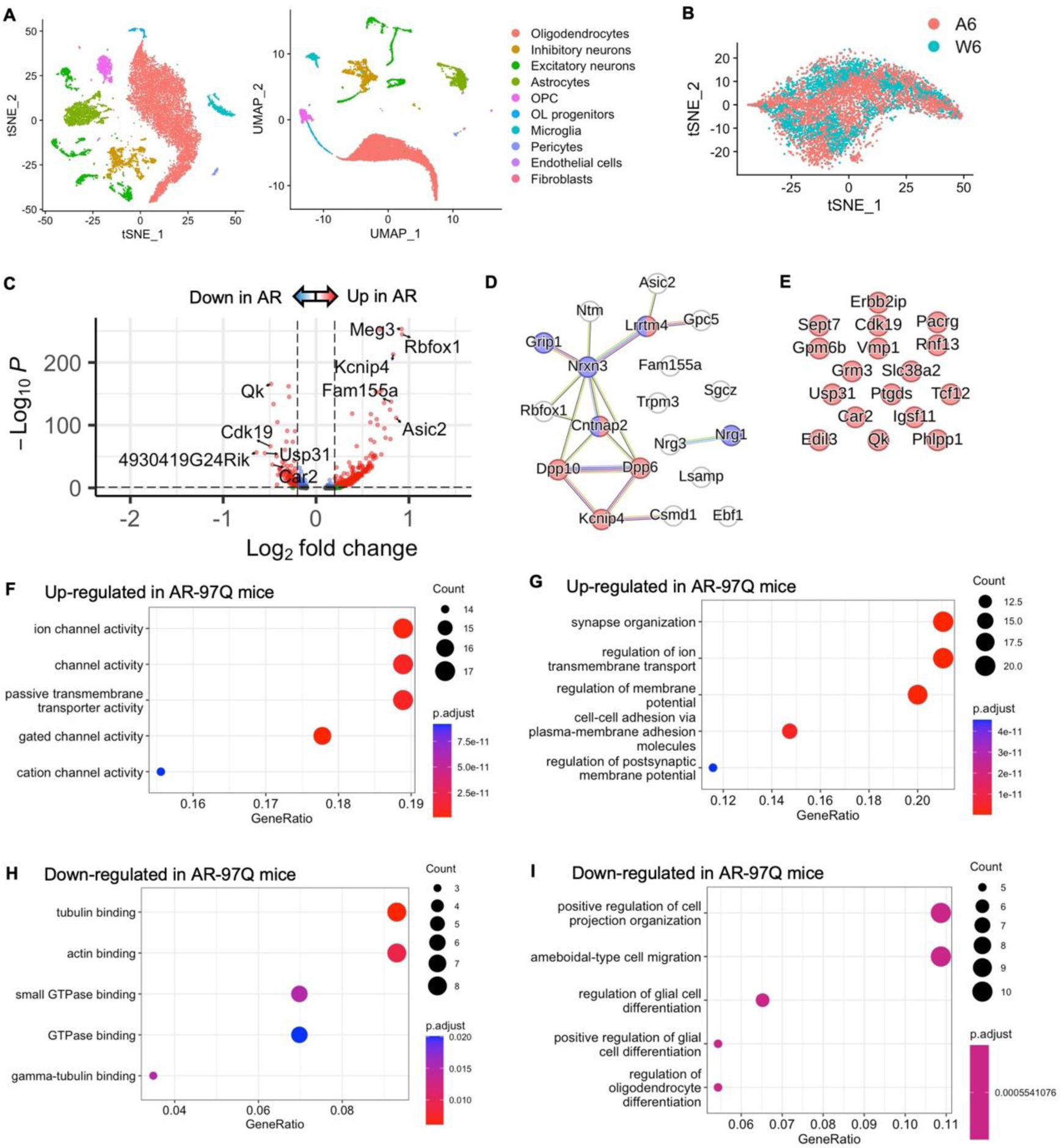
Unique gene expression signatures of the OL cluster at 6 weeks. **A,** t-Distributed stochastic neighbor embedding (t-SNE) and uniform manifold approximation and projection (UMAP) plots visualizing clusters of single nuclei in the spinal cord of AR-97Q and wild-type mice at 6 weeks. **B,** Genotype-colored t-SNE plot of the oligodendrocyte (OL) cluster: orange dots represent AR-97Q mice (A6), and green dots represent wild-type mice (W6). **C,** Volcano plot showing the DEGs of the OL cluster in AR-97Q mice and wild-type mice. The top 5 genes and last 5 genes are marked. **D,** Protein‒protein interaction (PPI) networks for the top 20 upregulated genes in AR-97Q mice. Genes colored in red have cation channel complex and genes colored in purple have synaptic membrane in the cellular component of GO terms. **E,** PPI networks for the top 20 downregulated genes in AR-97Q mice. **F, G,** The enrichment of the top 100 upregulated genes in AR-97Q mice in the biological process (**F**) and molecular function (**G**) categories (log_2_FC > 0.404). **H, I,** The enrichment of the top 100 downregulated genes in AR-97Q mice in the biological process (**H**) and molecular function (**I**) categories (log_2_FC < −0.224). A6, AR-97Q mice at 6 weeks; W6, wild-type mice at 6 weeks. Line color code: sky blue, known interactions from curated databases; magenta, experimentally determined interactions; green, predicted from neighborhood analysis; red, predicted from gene fusions; blue, predicted from gene cooccurrence; pastel green, text mining; black, coexpression; and clear violet, protein homology.

We reanalyzed microarray data from the whole spinal cords of AR-97Q mice at 7 to 9 weeks reported in a previous study (Minamiyama *et al*, 2012) (Supplementary Fig. 4A–C). A total of 25 genes were significantly upregulated in AR-97Q mice compared to AR-24Q mice (FDR < 0.1, FC > 1.5), and they were enriched in synapse assembly, suggesting that synaptic function is activated in the whole spinal cords of AR-97Q mice in the early stages of disease (Supplementary Fig. 4D). Microarray data also showed that the relative expression levels of genes related to cation channels and synaptic function were increased in the whole spinal cord of AR-97Q mice in the early stages of disease (Supplementary Fig. 4E, F), supporting our observation that they were among the top 20 upregulated genes in the OLs of AR-97Q mice at 6 weeks (Fig. 3D).

To elucidate the transcriptional changes in OLs after the onset of motor deficits, the data obtained from AR-97Q and wild-type mice at 9 weeks were compared, and 9 major cell types were specified (Supplementary Fig. 5A–C). The findings at 9 weeks were similar to those found at 6 weeks. The top 20 upregulated genes, but not downregulated genes in AR-97Q mice included several genes associated with the cation channel complex and synapses (Supplementary Fig. 5D, E). The top 100 upregulated genes in AR-97Q mice (log_2_FC > 0.338) were associated with ion channel activity and synapse organization (Supplementary Fig. 5F, G). The top 100 downregulated genes in AR-97Q mice (log_2_FC < −0.234) were related to actin binding, gliogenesis, and ensheathment of neurons (Supplementary Fig. 5H, I). Collectively, these findings suggest that axon sheath formation is impaired by 9 weeks of age.

To investigate OL heterogeneity in the advanced stages of SBMA, data from 13-week-old AR-97Q mice and wild-type mice were compared, and 9 major cell classes were identified (Fig. 4A–C). The top 20 downregulated genes showed interaction, but the top 20 upregulated genes in AR-97Q mice were predicted to be less connected to each other (Fig. 4D, E). The top 100 upregulated genes in AR-97Q mice (log_2_FC > 0.415) were associated with GTPase activator and regulator activity (Fig. 4F, G), although these genes were downregulated at 6 weeks (Fig. 3H). The top 100 downregulated genes in AR-97Q mice (log_2_FC < −0.436) were related to ion channel activity and synapse organization (Fig. 4H, I), although these pathways were activated at 6–9 weeks (Fig. 3F, G, Supplementary Fig. 5F, G). Altogether, our results demonstrated that ion channel activity and synapse organization were augmented in the early stages of disease but suppressed at the advanced disease stage.

**Fig. 4.**
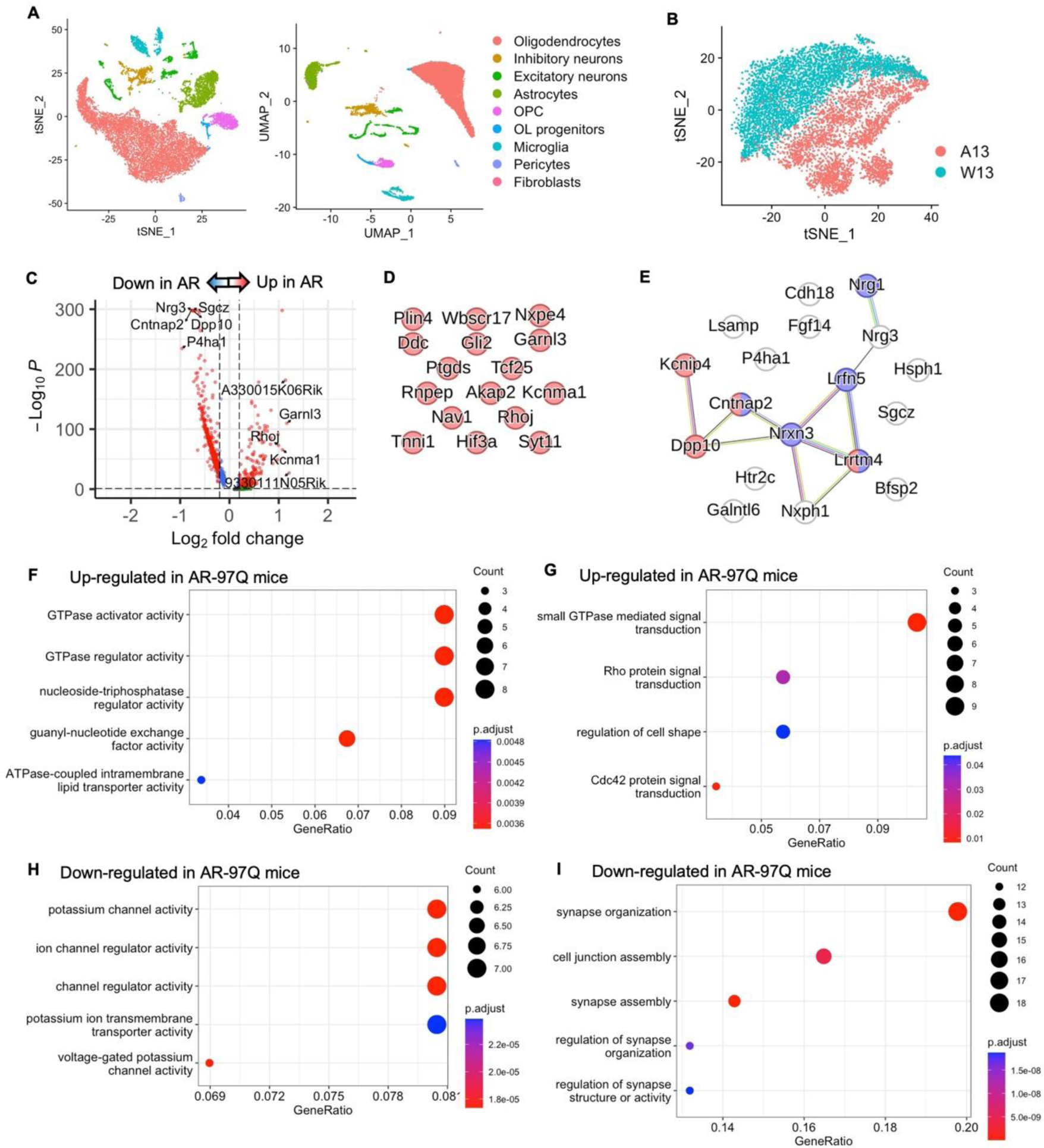
Unique gene expression signatures of the OL cluster at 13 weeks. **A,** t-Distributed stochastic neighbor embedding (t-SNE) and uniform manifold approximation and projection (UMAP) plots visualizing clusters of single nuclei in the spinal cord of AR-97Q and wild-type mice at 13 weeks. **B,** Genotype-colored t-SNE plot of the oligodendrocyte (OL) cluster: orange dots represent AR-97Q mice (A13), and green dots represent wild-type mice (W13). **C,** Volcano plot showing the DEGs in the OL cluster of AR-97Q mice and wild-type mice. The top 5 genes and last 5 genes are marked. **D,** Protein‒ protein interaction (PPI) networks for the top 20 upregulated genes in AR-97Q mice. **E,** PPI networks for the top 20 downregulated genes in AR-97Q mice. Genes colored in red have cation channel complex in the cellular component of GO terms and genes colored in purple have synaptic organization in the biological component of GO terms. **F, G,** The enrichment of the top 100 upregulated genes in AR-97Q mice in the biological process (**F**) and molecular function (**G**) categories (log_2_FC > 0.415). **H, I,** The enrichment of the top 100 downregulated genes in AR-97Q mice in the biological process (**H**) and molecular function (**I**) categories (log_2_FC < −0.436). A13, AR-97Q mice at 13 weeks; W13, wild-type mice at 13 weeks.

### Time course of differential gene expression

Three-week-old AR-97Q mice show neither motor symptoms nor nuclear aggregation of the polyglutamine-expanded AR (Fig. 2C), in agreement with the fact that prepubertal SBMA patients have no subjective symptoms (Atsuta *et al*, 2006). However, the toxicity of soluble polyglutamine oligomers, which appear before the onset of symptoms, has been shown in previous *in vivo* experiments (Li *et al*, 2007). We thus investigated the heterogeneity of OLs at 3 weeks by comparing the data of AR-97Q mice with those of wild-type mice at 3 weeks. Unsupervised clustering identified 8 major cell types, and the volcano plot of DEGs demonstrated fewer changes at 3 weeks of age compared to other ages (Supplementary Fig. 6A–C). A total of 17 upregulated genes in AR-97Q mice were predicted to be closely related to each other and involved several genes associated with the cation channel complex and synaptic membrane, but the interactions in downregulated genes were not clear (Supplementary Fig. 6D, E). The upregulated genes in AR-97Q mice (17 genes, log_2_FC > 0.22) were associated with ion channel activity and synapse organization (Supplementary Fig. 6F, G). The downregulated genes in AR-97Q mice at 3 weeks of age (67 genes, log_2_FC < −0.2) were related to small GTPase binding and carboxylic or organic acid biosynthetic processes (Supplementary Fig. 6H, I). The cluster of OLs at 3 weeks was reclustered into 6 subclusters for downstream analysis (Supplementary Fig. 7A, B). Based on the cell subpopulation proportional diagram, subcluster 0 was mainly enriched in AR-97Q mice and was associated with neurexins and neuroligins, protein‒protein interaction at synapses, and the neuronal system according to Reactome pathway analysis (Supplementary Fig. 7C, D). Altogether, the transcriptional changes observed in AR-97Q mice at 6 and 9 weeks were also observed to a lesser extent at 3 weeks.

To understand the mechanisms of OL transcriptional changes in the absence of nuclear aggregation, we searched ChIP-Atlas (http://chip-atlas.org) for transcription factors that are associated with the top 10 and last 10 DEGs in OLs at 3 weeks (Supplementary Fig. 8). Several of these transcription factors have been reported to interact with AR, suggesting that these genes are involved in the transcriptional dysregulation of *AR* even before nuclear aggregation occurs.

To investigate the timeline of transcriptional changes in SBMA, data from AR-97Q mice in four disease stages were compared (Fig. 5A, B). The top 100 upregulated genes in the OLs of AR-97Q mice at 13 weeks (log_2_FC > 0.275) compared with those at 3, 6, and 9 weeks were associated with small GTPase-mediated signal transduction activators (Fig. 5C). In contrast, the top 100 downregulated DEGs in AR-97Q mice at 13 weeks (log_2_FC < −0.100) were related to synapse organization and cell junction assembly (Fig. 5D). These findings were consistent with the DEGs between the two groups at each week. We utilized pseudotime trajectory analysis to reveal the differentiation process of OL lineage cells. The results showed that they formed a continuous differentiation trajectory from OPCs to OL progenitors to OLs (Fig. 5E, F). The boxplot shows that cells at 13 weeks exhibited a different pseudotime pattern from those at 3, 6, and 9 weeks (Fig. 5G). The pseudotime kinetics of *Pdgfrα* (a marker of OPC), *Sox6* (a marker of OPCs and OL progenitors), and *Mog* (a marker of OLs) revealed that cluster 8 represented OPCs, cluster 20 represented OL progenitors, and other clusters represented OLs (Fig. 5H–L). Compared to other OL clusters, cluster 4, the predominant cluster at 13 weeks, exhibited high expression of *Tnr*, a marker of OPCs and OL progenitor cells, and low expression of *Apc*, a marker of mature OLs, indicating that OLs at 13 weeks are immature (Fig. 5M, N). In contrast, the OL lineage cells of wild-type mice showed almost no deviation in pseudotime distribution over time compared to those of AR-97Q mice (Supplementary Fig. 9).

**Fig. 5.**
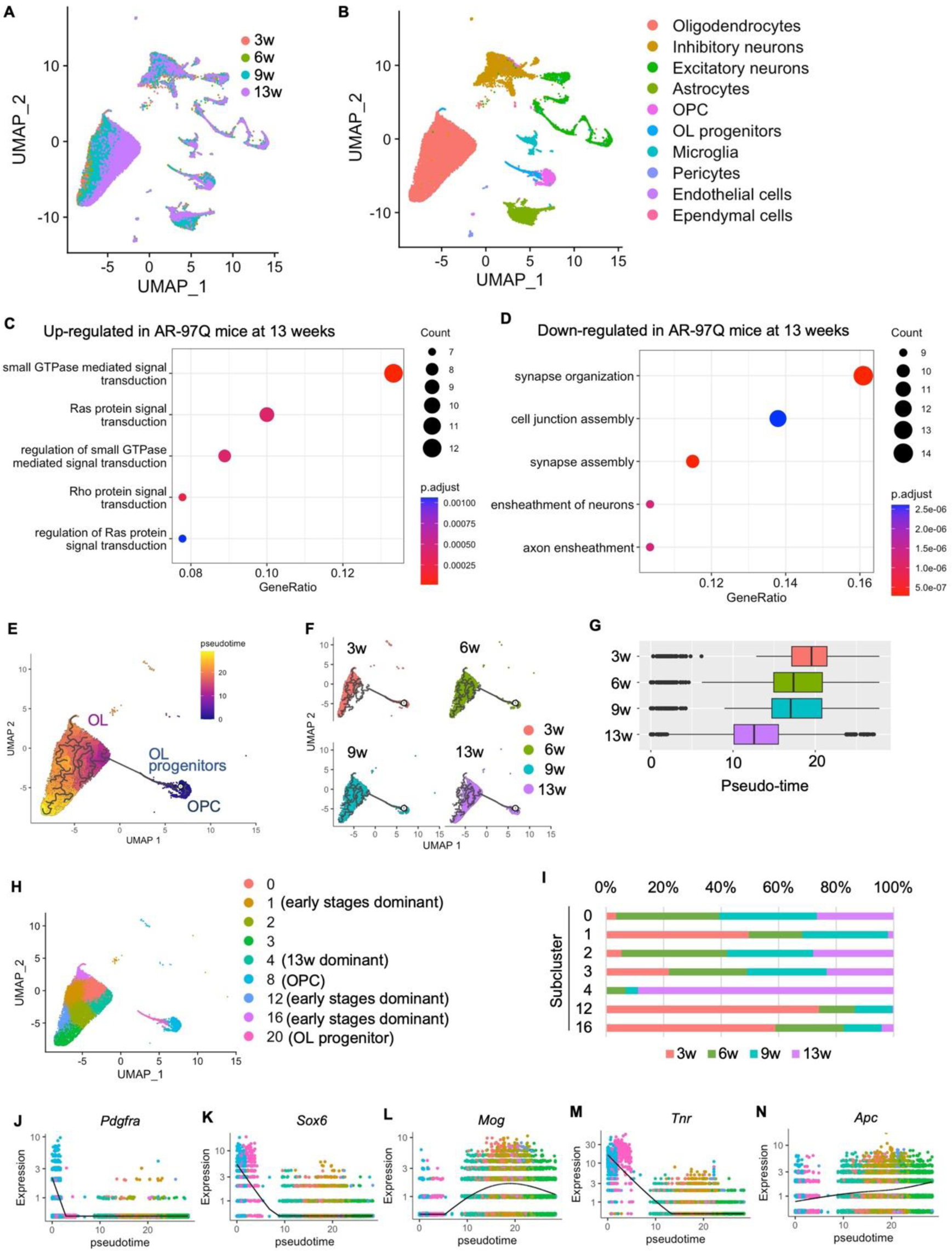
Transcriptional changes in OL according to disease stage. **A,** Uniform manifold approximation and projection (UMAP) plots of all AR-97Q mice samples color-coded by each week of age (resolution = 1.2). **B,** UMAP plots of each cell type of AR-97Q mice at four disease stages. **C, D,** The enrichment of 100 genes upregulated (**C**) or downregulated (**D**) in AR-97Q mice at 13 weeks in the biological processes category (log_2_FC > 0.275 or log_2_FC < −0.1). **E,** Pseudotime analysis inferred from the OL lineage cell clusters of AR-97Q mice in 4 disease stages. **F,** UMAP visualization of OL lineage cells clusters colored by weeks of age. **G,** Boxplot showing the distribution of pseudotime within each sample. Vertical bars indicate median values. **H,** UMAP visualization of OL lineage cell clusters colored by the Seurat package. **I,** Proportion of each subcluster of each sample. **J-N,** Pseudotime kinetics of *Pdgfra* (**J**), *Sox6* (**K**), *Mog* (**L**), *Tnr* (**M**), and *Apc* (**N**).

### The levels of ion channel-related proteins are increased in the early stages of SBMA

The top 20 upregulated genes in AR-97Q mice at each week were compared to elucidate the broad range of changes that are common in the early disease stages (Fig. 6A). *Asic2*, *Fam155a*, *Meg3*, and *Rbfox3* were among the top 20 shared DEGs at 3, 6 and 9 weeks of age. Among them, the expression levels of *Asic2* and *Fam155a*, which are associated with cation channels, specifically sodium channels, in the OLs of AR-97Q mice were increased more than 1.75-fold at 6 weeks and less than 0.72-fold at 13 weeks compared to those in the OLs of wild-type mice (Fig. 6B–D). The pseudotime analysis of *Asic2* and *Fam155a* showed that their expression levels were elevated in clusters 1, 12, and 16, early stage-dominant clusters, and decreased in cluster 4, a late stage-dominant cluster (Fig. 5I, 6E, F). The analysis of OL lineage cells of wild-type mice showed almost no deviation in the expression levels of *Asic2* and *Fam155a* among the four stages (Supplementary Fig. 9, 10). Immunofluorescence analysis of AR-97Q mouse spinal cords confirmed the increased levels of Asic2 and Fam155a in OLs at 6 weeks and their decreased levels in OLs at 13 weeks compared to those in the OLs of wild-type mice (Supplementary Fig. 11, 12).

**Fig. 6.**
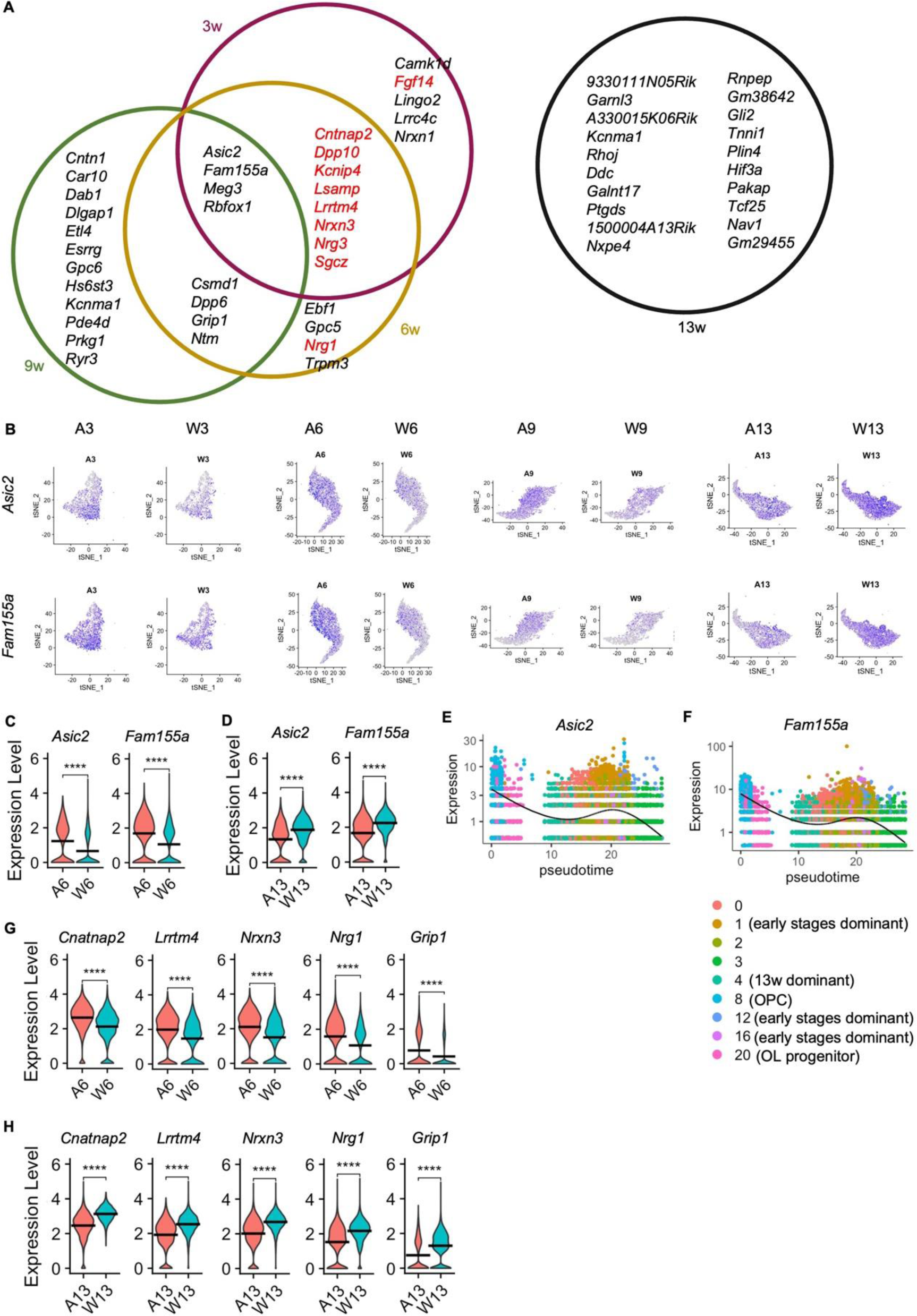
Top 20 DEGs in the OLs of AR-97Q mice and wild-type mice in each week. **A,** Common top 20 DEGs across the different weeks of age. Genes in red are the top 20 genes that are downregulated in AR-97Q mice at 13 weeks. **B,** Feature plots of *Asic2* and *Fam155a* expression in the OL cluster of each sample. **C, D,** Violin plots of *Asic2* and *Fam155a* expression in the OLs of AR-97Q mice and wild-type mice at 6 weeks (**C**) and 13 weeks (**D**). **E, F,** Pseudotime kinetics of *Asic2* (**E**) and *Fam155a* (**F**) in the OL lineage cells of AR-97Q mice at 3, 6, 9, and 13 weeks. UMAP visualization of OL lineage cell clusters in AR-97Q mice is shown in Fig. 5G**. G, H**, Violin plots of *Cntnap2*, *Lrrtm4*, *Nrxn3*, *Nrg1*, and *Grip1* expression in the OLs of AR-97Q mice and wild-type mice at 6 weeks (**G**) and 13 weeks (**H**). Vertical bars indicate mean values (**C, D, G, H**). \*\*\*\**p* < 0.0001.

### Genes associated with synaptic activity are upregulated in the early stages of SBMA

The top 20 upregulated genes in the OLs of AR-97Q mice at 3 and/or 6 weeks included 10 of the top 20 genes that were downregulated in the OLs of AR-97Q mice at 13 weeks (Fig. 6A). Such genes include *Cntnap2*, *Lrrtm4*, *Nrxn3*, and *Nrg1*, and the common enriched GO cellular component term of these genes was the synaptic membrane (Fig. 6G, H). Moreover, proteins synthesized from three of these genes, Cntnap2, Lrrtm4, and Nrxn3, have been reported to be coexpressed (Fig. 3D, 4E, Supplementary Fig. 6D). Additionally, cation channel complexes was an enriched GO term of *Cntnap2* and *Lrrtm4*. Cntnap2 is needed for the normal clustering of potassium channels, and Lrrtm4 interacts with Nrxn in a calcium-dependent manner. Four genes, *Csmd1*, *Dpp6*, *Grip1*, and *Ntm*, were among the top 20 upregulated genes in the OLs of AR-97Q mice at 6 and 9 weeks (Fig. 6A), and they were significantly downregulated in AR-97Q mice at 13 weeks. Grip1 and Ntm localize to the synapse, and specifically, *Grip1* is the 36th DEG that was downregulated in the OLs of AR-97Q mice at 13 weeks (Fig. 6G, H). Grip1 interacts with GluR2, a common subunit of the α-amino-3-hydroxy-5-methyl-4-isoxazolepropionic acid receptor (AMPAR), and plays a critical role in delivering AMPARs to excitatory synapses.

### Transcriptional dysregulation in other cell types

To further understand the dysregulation of other cell types in SBMA, the upregulated genes among the OLs, astrocytes, and microglia clusters and the OLs, inhibitory neurons, and excitatory neurons clusters in AR-97Q mice were compared (Supplementary Fig. 13). *Camk1d* was commonly upregulated in the OLs, astrocytes, inhibitory neurons, and excitatory neurons clusters at 3 weeks. At 6 weeks, 6 genes were universally upregulated in OLs, astrocytes, and microglia clusters in AR-97Q mice, and three of them, *Asic2*, *Meg3*, and *Rbfox1*, were among the top 20 upregulated genes in OLs throughout 3, 6, and 9 weeks of age. The top 20 DEGs had nothing in common with OLs, inhibitory neurons, and excitatory neurons at 6 weeks.

We also explored the transcriptional changes in the OPCs of AR-97Q mice, because the OLs exhibited changes from the very beginning of disease (Supplementary Fig. 14). The upregulated genes in the OPCs of AR-97Q mice at 6 weeks (18 genes, log_2_FC < 0.37) were associated with synapse assembly and cell junction assembly. At 13 weeks, the top 100 downregulated genes in the OPCs of AR-97Q mice (log_2_FC < −0.368) were associated with ion transmembrane transport and cell junction assembly. Changes in the OPCs of AR-97Q mice were similar to those in the OLs of AR-97Q mice, indicating that the alterations in OLs are similar across OL lineage cells.

### The SBMA OL cell model reflects the early pathogenesis of SBMA

To further validate the transcriptional alterations identified in snRNA-seq analysis, we generated an oligodendroglial cell model of SBMA using Oli-neu cells (Fig. 7A). Cell differentiation was induced by adding PD174265, a selective inhibitor of the EGF receptor that has been previously shown to be a differentiating agent for Oli-neu cells, and was confirmed by increased levels of Mog and decreased expression levels of Pdgfrα (Naffaa *et al*, 2022) (Fig. 7B). Total RNA from differentiated AR-17Q cells and AR-97Q cells was isolated, and gene expression analysis was performed. Eleven genes were significantly upregulated (> 2-fold), and 86 genes (less than one-half) were significantly downregulated in AR-97Q cells (p < 0.05), and hierarchical clustering analysis revealed a clear difference between AR-17Q cells and AR-97Q cells (Fig. 7C–E). The DEGs were associated with the regulation of signaling and cell communication, and the synaptic membrane was the most enriched GO cellular component term (Fig. 7F, G), indicating that the DEGs may function in synapse or signal transduction and that the OL cell model of SBMA showed phenotypes similar to those of AR-97Q mice. To compare the OLs in the SBMA cell model with those in AR-97Q mice, genes that were upregulated or downregulated in both the cell and mouse models of SBMA were selected (|log_2_FC| > 0.1, adjusted p value < 0.05). The ratio of the number of these genes to the number of genes for which the expression levels were commonly measured by both RNA-seq methods was calculated. The results revealed that the ratio was significantly lower at 13 weeks than at other weeks, suggesting that the OLs in the SBMA cell model were more similar to those in AR-97Q mice in early disease stages than in advanced stages (Supplementary Fig. 15A). The top 6 enriched GO terms of the genes that were upregulated in the OLs of both the cell model and AR-97Q mice (log_2_FC > 0.1) at 6 weeks involved postsynapse organization and action potential, indicating that genes related to synapses are upregulated in both the SBMA cell and animal model OLs (Supplementary Fig. 15B). Quantitative real-time polymerase chain reaction (RT‒PCR) analyses showed that the mRNA levels of *Asic2*, *Fam155a*, *Cntnap2*, and *Grip1* in AR-97Q cells were elevated, indicating that the genes related to ion channels and synapse function were also increased in the OLs of the SBMA cell model as well as in the OLs of AR-97Q mice in early disease stages (Fig. 6C, G, Fig. 7H).

**Fig. 7.**
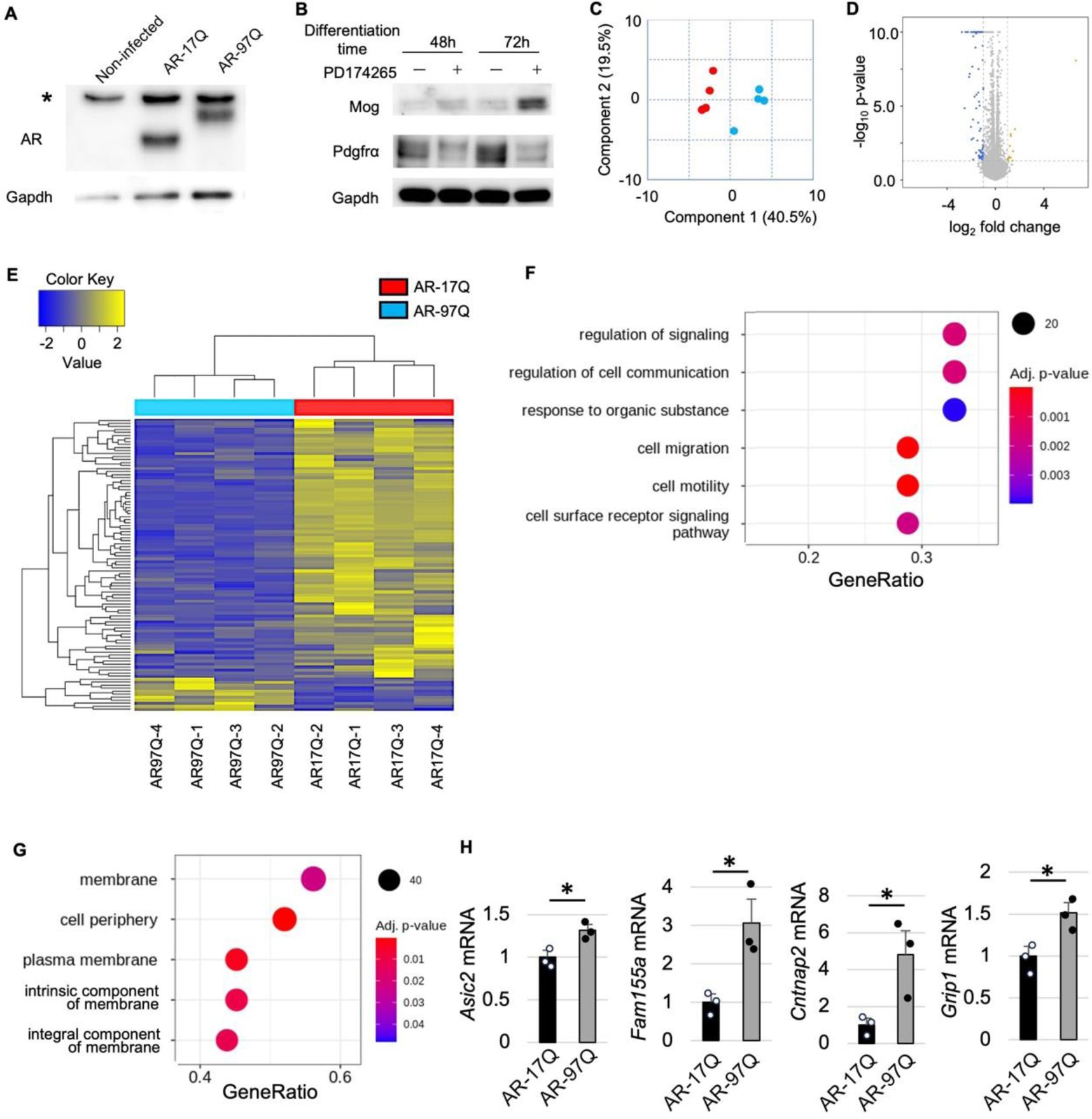
OL cell model of SBMA using the Oli-neu cell line. **A,** Immunoblots for human AR in noninfected, AR-17Q, and AR-97Q cells. **B,** Immunoblots showing the expression levels of Mog and Pdgfrα in AR-97Q cells with or without PD174265. **C,** Multidimensional scaling analysis of samples submitted to RNA-seq. Blue dots indicate samples of AR-97Q cells, and red dots indicate samples of AR-17Q cells. **D,** Volcano plot of the expression levels of the two groups. **E,** Heatmap showing the results of hierarchical clustering analysis, which clusters genes based on their significant DEGs. **F, G,** GO terms related to the biological process (**F**) and cellular component (**G**) that are enriched in the DEGs identified by RNA-seq. **H,** The mRNA levels of *Asic2*, *Fam155a*, *Cntnap2*, and *Grip1* in AR-17Q cells and AR-97Q cells. N = 3 samples for each group. Error bars indicate the SEM. \**p* < 0.05, unpaired two-sided t test. *, nonspecific bands.

### Interaction strength between OLs and excitatory neurons is elevated in the early stages of SBMA

Finally, we evaluated cell‒cell communication patterns in the spinal cords of AR-97Q and wild-type mice by applying CellChat to the scRNA-seq dataset. A heatmap of the differential number of interactions and differential interaction strength in AR-97Q mice compared to wild-type mice showed that the number of interactions and interaction strength between cells were elevated overall at 6 and 9 weeks and suppressed at 13 weeks (Fig. 8A–C). Similarly, when focusing on OLs, the number of interactions with other cell types, including neurons, were increased at 6 and 9 weeks and decreased at 13 weeks. Analysis of OL signaling in AR-97Q mice compared to wild-type mice revealed that either the incoming or outgoing interaction strength of the NRG, NRXN, and NCAM pathways, which are associated with neuronal function, was increased in AR-97Q mice at 6 and 9 weeks (Fig. 8D, E). At 13 weeks, the incoming and outgoing interaction strength of the NRXN pathway was elevated in the OLs of AR-97Q mice (Fig. 8F). Signaling changes in the OL progenitors and OPCs of AR-97Q mice compared to wild-type mice showed similar trends to the signaling changes in OLs. To clarify the effects of OLs on inhibitory neurons and excitatory neurons, the communication probability of the significant ligand‒receptor pair interactions between OLs and inhibitory neurons or excitatory neurons was calculated. NCAM and NRXN signaling was highly elevated between the OLs and inhibitory neurons or excitatory neurons at 6 and 9 weeks (Fig. 8G, H), and it was decreased at 13 weeks (Fig. 8I). Ptn, Negr1, Lrrc4c, Efnb3, and Cadm1 signaling was also increased between OLs and neurons at 6 and 9 weeks, and it was decreased at 13 weeks. Many of these factors are associated with cell adhesion and synapse formation.

**Fig. 8.**
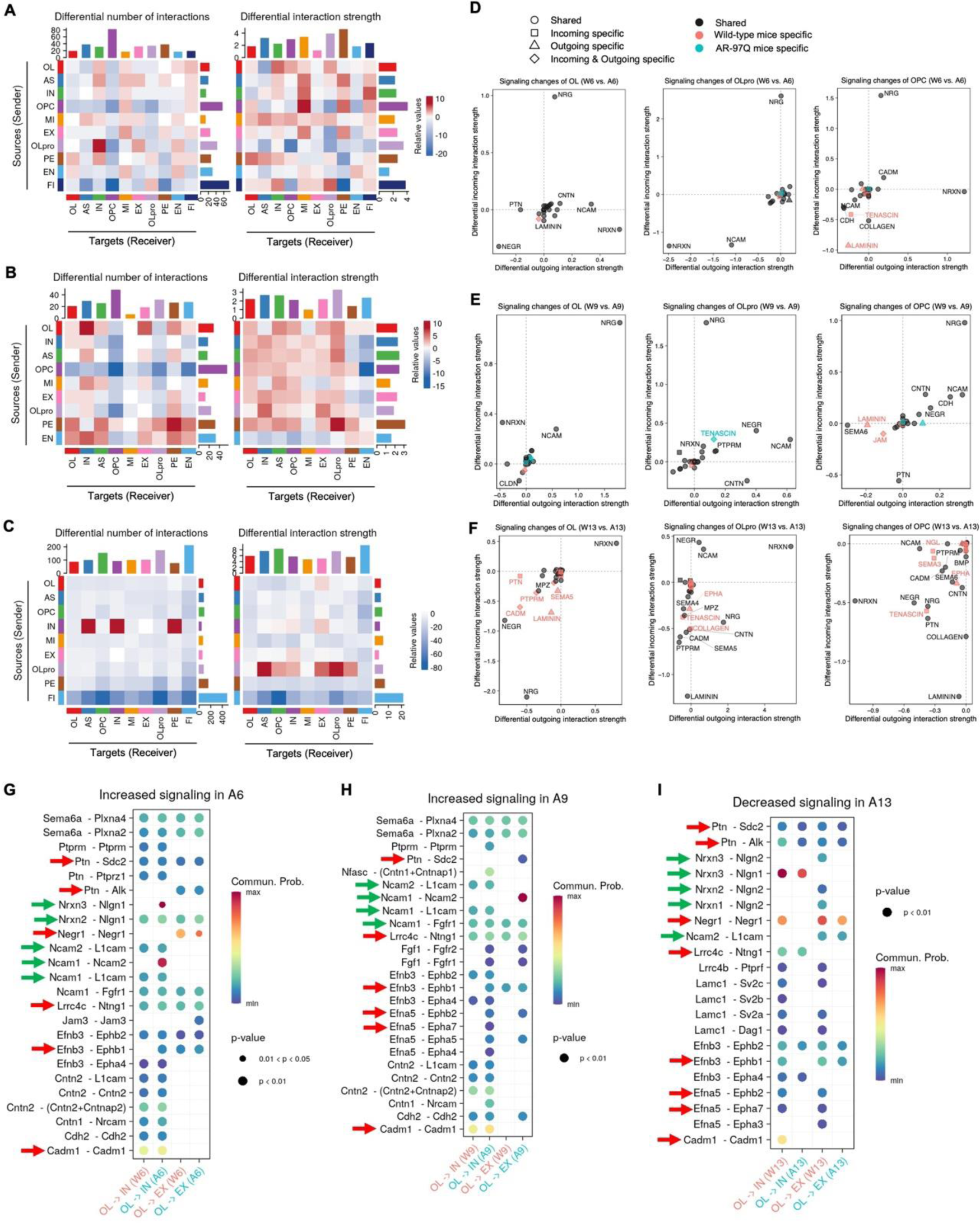
Inferred intercellular communication network in the spinal cords of AR-97Q mice. **A–C,** Heatmap of differential interaction strength in AR-97Q mice compared to wild-type mice at 6 weeks (**A**), 9 weeks (**B**), and 13 weeks (**C**). The top colored bar plot represents the sum of the values in columns displayed in the heatmap (incoming signaling). The right colored bar plot represents the sum of the values in rows (outgoing signaling). In the heatmap, red (or blue) represents increased (or decreased) signaling in AR-97Q mice compared to wild-type mice. Relative value = the interaction strength from source to target in AR-97Q mice – the interaction strength from source to target in wild-type mice. **D–F,** Signaling changes in OLs, OL progenitors, and oligodendrocyte precursor cells (OPCs) in AR-97Q mice compared to wild-type mice at 6 weeks (**D**), 9 weeks (**E**), and 13 weeks (**F**). **G–I,** Bubble plots of the communication probability of all the significant ligand‒receptor interactions between OLs and inhibitory neurons or excitatory neurons, which are increased in AR-97Q mice at 6 weeks (**G**) and 9 weeks (**H**) and decreased in AR-97Q mice at 13 weeks (**I**). The dot color and size represent the communication probability and p values, respectively. The p values were computed from a one-sided permutation test. The ligand‒ receptor pair interactions that are increased in AR-97Q mice at 6 and 9 weeks and decreased at 13 weeks are indicated by red arrows. The ligand‒receptor pairs related to NCAM and NRXN are indicated by green arrows. AS, astrocytes; IN, inhibitory neurons; MI, microglia; EX, excitatory neurons; OLpro, OL progenitors; PE, pericytes; EN, endothelial cells; FI, fibroblasts.

## Discussion

In the present study, we identified cell type- and disease stage-specific gene expression changes in the spinal cord of SBMA model mice using snRNA-seq analysis. The transcriptional changes in OLs were the most evident among all cell types; the number of DEGs in OLs was the highest among all cell types in the early stages of disease. GO and pathway analyses of the DEGs in the OL clusters of SBMA model mice at each week showed that pathways associated with ion channels and synapses were activated in the OLs of AR-97Q mice from the preonset to early symptomatic stage of disease, suggesting that cell hyperexcitation and the reinforcement of intercellular connections occur early in SBMA. OLs are myelinating cells in the central nervous system that mediate the rapid conduction of action potentials and provide trophic support for axonal and neuronal maintenance. The supply of energy metabolites from OLs to axons is crucial in the neurological system. Disease-associated OL signatures have emerged as important contributors to the development of neurodegenerative diseases such as Huntington’s disease (HD) (Yi Teo *et al*, 2016; Huang *et al*, 2015; Lim *et al*, 2022; Labadorf *et al*, 2015; Wilson *et al*, 2018), Parkinson’s disease (PD) (Bryois *et al*, 2020), and Alzheimer’s disease (AD) (Maitre *et al*, 2023; Ferrer & Andrés-Benito, 2020); however, the role of OLs in the pathogenesis of SBMA has yet to be elucidated. OLs and OPCs create functional unidirectional or bidirectional synaptic contacts and regulate synaptic plasticity (Bradl & Lassmann, 2010; Lin & Bergles, 2004; Zhang *et al*, 2021). Myelin-forming OLs detect electrical activity in axons and increase intracellular calcium levels (Micu *et al*, 2007), and alternatively, the depolarization of OLs further increases the conduction velocity of myelin-forming axons (Yamazaki *et al*, 2007). It was also demonstrated that OL depolarization increases the number of excitatory synaptic responses and facilitates the induction of long-term potentiation at synapses (Yamazaki *et al*, 2019).

In the present study, genes related to ion channel activity were upregulated in the OL clusters of AR-97Q mice at 3, 6, and 9 weeks, suggesting that depolarization and repolarization are abnormally activated at the early stage of SBMA. Moreover, Asic2, an acid-sensing ion channel 2, is known to be localized at the synapse and promote long-term potentiation and synaptic plasticity. Recent studies on the pathophysiology of multiple sclerosis (MS) have indicated that the concentrations of calcium and sodium ions affect the degree of myelin damage. Analysis of human autopsied brain tissue showed increased *ASIC2* mRNA levels in MS samples compared to control subject samples (Fazia *et al*, 2019).

Neurons overexpressing *ASIC2a* fired more frequently than control neurons (Zhang *et al*, 2017), suggesting the involvement of ASIC2 in the neuroexcitatory imbalances observed in epilepsy. Fam155a, also known as NALCN channel auxiliary factor 1 (NALF1), is a voltage-gated ion channel responsible for the resting sodium permeability, which controls neuronal excitability. Ion channels are critical components of cellular excitability, and cell hyperexcitation has been implicated in the pathogenesis of several neurodegenerative diseases. It has been suggested that hyperexcitability in amyotrophic lateral sclerosis (ALS) is driven by changes in voltage-gated sodium and potassium channels (Kanai *et al*, 2012), as shown in postnatal ALS model mice, motor neurons cultured from ALS model mice, and induced pluripotent stem cells (iPSCs) with mutant *SOD1*, *C9Orf72*, or *FUS* (Wainger *et al*, 2014; Van Zundert *et al*, 2008). Studies on ALS patients have shown that persistent sodium conductance is strongly associated with shorter survival, suggesting that alterations in cell excitability are pathogenic (Kanai *et al*, 2012). Altogether, our results suggest that the sodium current is disrupted by OLs, which is in line with the observed benefit of mexiletine, a sodium channel blocker, in patients with SBMA (Yamada *et al*, 2022).

We also found that genes associated with synaptic function were upregulated in the early stages of SBMA, whereas they were downregulated in the advanced stage. In particular, among the top 20 DEGs at 3 and/or 6 weeks of age, Cntnap2, Lrrtm4, Nrxn3, and Nrg1 are known to localize at the synaptic membrane, mediate cell‒cell signaling, and regulate synaptic plasticity. Disruption of Cntnap2 level leads to imbalanced excitatory and inhibitory neural networks, which are thought to be central to the pathophysiology of schizophrenia (St George-Hyslop *et al*, 2023). Lrrtm4 is a postsynaptic protein that regulates the development and strength of glutamatergic synapses by interacting with presynaptic proteins such as Nrxns (Sakamoto *et al*, 2021). Increased or decreased levels of Nrg1 result in the abnormal growth of dendritic spines and the disruption of excitatory and inhibitory synapses, suggesting that maintaining a balance of synaptic protein levels is critical to preserve synaptic function. Among the top 20 DEGs at 6 and 9 weeks of age, Grip1 plays an essential role in regulating AMPAR trafficking during synaptic plasticity and learning and memory (Tan *et al*, 2020). The expression and synaptic distribution of Grip1 is disturbed in the pathological state, resulting in the increased susceptibility of neurons to AMPA-induced neurotoxicity (Lai *et al*, 2006). *Cntnap2*, *Dpp10*, *Kcnip4,* and *Dpp6* are involved in the potassium channel complex, and potassium channels are key regulators of synaptic plasticity (Voglis & Tavernarakis, 2006). Ion channels at synapses contribute to proper synaptic function by regulating membrane potential and participating in neurotransmitter release and vesicle recycling. This close relationship between channels and synapses resulted in transcriptional changes in the present study, with both ion channel and synaptic genes being upregulated in early stages of the disease and downregulated in advanced stages. Collectively, our results indicate deficits in ion channel and synaptic functioning in OLs in the early stages of SBMA.

The stage-specific changes in cell activities have been reported in other neurodegenerative diseases. Clinical studies using functional magnetic resonance imaging have shown cortical and hippocampal hyperactivity in the early stages of AD, progressing to hypoactivity in the later stages of neurodegeneration (Targa Dias Anastacio *et al*, 2022; Celone *et al*, 2006; Dickerson *et al*, 2005). A previous study reported a greater number of hyperactive CA1 pyramidal neurons in the hippocampus of AD model mice than wild-type mice at an early disease stage; however, this number was reduced as the mice aged (Busche *et al*, 2012). Hyperexcitability occurs in the early stages of ALS, even prior to motor symptom onset, and then progresses to hypoactivity in later stages (Do-Ha *et al*, 2018). In an analysis using trans-magnetic stimulation, patients in the early stages of ALS showed a hyperexcitable motor cortex, whereas patients in the later stages of ALS exhibited a hypoexcitable motor cortex compared to the control group (Vucic & Kiernan, 2006). Transcriptional changes in the early stages of SBMA observed in the current study may reflect the pathological mechanisms underlying the neuromuscular degeneration characteristic of SBMA, while transcriptional changes at 13 weeks may indicate consequential and/or compensatory changes after neurodegeneration.

Intercellular network analysis revealed changes in the interactions between OLs and neurons and allowed us to infer a mechanism by which OL abnormalities drive neurodegeneration. Nrxn3 signaling between OLs and inhibitory neurons was significantly elevated at 6 weeks, which is consistent with the snRNA-seq results showing that *Nrxn3* was among the top 20 DEGs in the OL cluster of AR-97Q mice in the early stages of disease. Moreover, Ncam1–Ncam2 signaling between OLs to inhibitory and excitatory neurons was inferred to be increased in the early disease stages. Elevated NCAM2 levels and altered submembrane Ca^2+^ dynamics are known to cause defects in synapse maturation, as implicated in the pathology of Down syndrome and other brain disorders associated with abnormal NCAM2 expression (Sheng *et al*, 2019). The increased strength of NRG, NRXN, and NCAM signaling between OLs and other cell types can lead to abnormalities in synaptic function as well as abnormalities in the function of the cell receiving the signal.

This study has several limitations. First, we confirmed the gene expression changes in OLs, but the function of ion channels and synapses should be verified in the future by cell coculture or animal experiments. Second, although we demonstrated OL dysfunction in autopsied patient tissues, early changes in SBMA patients remain elusive. Development of biomarkers that can be measured in biosamples of patients and carriers of SBMA is needed.

## Materials and Methods

### Animals

AR-97Q (Line #7–8) mice were bred and maintained in the animal laboratory of our institute (Katsuno *et al*, 2002). The mice were genotyped by PCR amplification using DNA extracted from the tail with the primers listed in Supplementary Table 1.

### Tissue processing for single-nucleus sequencing

C57BL6 and AR-97Q male mice were deeply anesthetized, the spinal column was cut at the hip level, and a PBS-filled syringe fitted with a 20 G needle (7.25 mm) was inserted into the caudal end of the spinal column to flush out the spinal cord. Spinal cords were snap-frozen in powdered CO_2_ in liquid nitrogen and stored at −80°C until processing. Spinal cords from 4 mice per group were pooled for homogenization and nuclear isolation. The nuclear isolation protocol was adapted from the ‘Frankenstein’ protocol. Each sample was thawed and homogenized in a Dounce Homogenizer (Kimble Chase 2 ml Tissue Grinder) containing 500 μl freshly prepared ice-cold lysis buffer (10 mM Tris-HCl pH 7.4, 10 mM NaCl, 3 mM MgCl2, 0.1% Igepal and 0.2 U/μl SUPERase-In^TM^ RNase Inhibitor (#AM2696, Thermo Fisher Scientific, Waltham, MA, USA)). Then, 900 μl of lysis buffer was added, and the samples were incubated for 5 minutes. The homogenate was filtered through a 70 µm cell strainer (#08-771-2, Thermo Fisher Scientific) and centrifuged at 500×g for 5 min at 4 °C. The supernatant was removed, and the pellet was gently resuspended in 1.5 ml of lysis buffer and incubated for another 3 minutes on ice. Then, the nuclei were centrifuged at 500×g for another 5 min and resuspended in 1500 µl of wash buffer (1× PBS with 1% BSA and 0.2 U/µl SUPERase-In^TM^ RNase Inhibitor). The nuclei were washed and centrifuged at 500 × g for 5 minutes three times and filtered through a 40 µm cell strainer (#08-771-1, Thermo Fisher Scientific). Then, Debris Removal Solution (#130-109-398, Miltenyi Biotec Inc. CA, USA) was added to the cell suspension to create a density gradient and remove dead cells and debris. The samples were resuspended in 3.1 ml of 1x PBS in a 15 ml tube, and 900 μl of Debris Removal Solution was added and mixed well. The solution was gently overlaid with 4 ml of cold 1X PBS. The sample was centrifuged at 1000×g for 10 min at 4°C with fast acceleration and braking. The top two layers were aspirated and discarded. The bottom layer was left undisturbed, and the volume was increased to 15 ml with 1X PBS. Cells were gently mixed and centrifuged at 1000 g for 10 min at 4°C. The supernatant was removed, and the cells were resuspended in cold 0.04% BSA/PBS and stained with DAPI. Finally, nuclei labeled with DAPI were sorted with a FACS Aria instrument (Becton Dickson, MD, USA) with a 100 nm nozzle and 405 nm excitation laser. The instrument was controlled by a PC running FACS DiVa™ software (Becton Dickson). We collected at least 500,000 DAPI-positive nuclei from each condition. These nuclei were centrifuged and inspected for visual appearance, and the concentration was adjusted to proceed to the next step.

### Single-nucleus RNA sequencing

A total of 10,000 nuclei per sample were run on the 10× Chromium Single Cell 3’ gene expression v3.1 platform. DAPI-positive nuclei were immediately loaded onto a Chromium Single Cell Processor (10X Genomics) to barcode the mRNA inside each nucleus. Sequencing libraries were constructed according to the manufacturer’s instructions, and the resulting cDNA samples were run on an Agilent Bioanalyzer using the High Sensitivity DNA Chip as a quality control to determine cDNA concentrations. The samples were run on an Illumina HiSeq X Ten with Read 1 = 28 bp and Read 2 = 90 bp to obtain >= 20K reads per cell. A total of 70,046 cells with 1.96 billion reads were sequenced for the 8 samples with an average of 8756 cells per sample with 28.7 K reads each. Reads were aligned and assigned to Ensembl mm10 transcript definitions using the CellRanger v5.0.1 pipeline (10X Genomics). The gene barcode matrices for each sample were imported into R using the Read10X function in the Seurat R package (v3.1.460) (Hao *et al*, 2021).

### Quality control and filtering

Based on the distribution of the number of genes detected in each cell and the distribution of the number of unique molecular identifiers (UMIs), nuclei with fewer than 200 genes or more than 3000 genes were excluded from the downstream analyses. Nuclei with more than 1% of mitochondrial gene expression were excluded. Removal of outliers resulted in 54,456 total remaining cells for analysis. UMI counts were then normalized in Seurat 3.0, and the top 3000 highly variable genes were identified using the FindVariableFeatures function with variance stabilization transformation (VST).

### Dimension reduction and cluster annotation

We performed a reference-based integration workflow with reciprocal principal component analysis. Clustering was performed using the Seurat functions FindNeighbors and FindClusters. Visualization of the integrated dataset was performed using t-SNE and UMAP, with the first 25 principal components at a resolution of 0.8 if not otherwise noted. For subclustering, OL clusters were taken as a subset from the integrated dataset of wild-type and AR-97Q mice at 3 weeks, and significant PCs were used for downstream clustering similar to above. The cell type designation was established by first analyzing the DEGs in each cluster and manually comparing them to several canonical markers for each cell type.

### Differential expression analysis and GO enrichment analysis

Differential expression analysis between the two conditions was performed using FindMarkers. For the identification of DEGs, the log fold-change mentioned in each test and Wilcoxon’s signed rank test with Bonferroni correction for p value (< 0.05) were used as cutoff values. GO enrichment analysis and Reactome pathway analysis were performed to determine the function of the DEGs using the R package clusterProfiler (Yu *et al*, 2012).

### Pseudotime analysis of cell differentiation trajectories

The Monocle3 package (v1.3.1) (Trapnell *et al*, 2014) was used for pseudotime trajectory analysis to determine the differentiation status of cells at different disease stages.

### Cell‒cell interaction analysis

Seurat preprocessed data were subjected to the CellChat package (v1.6.1) with the Seurat R package (v4.3.1) to infer, analyze, and visualize cell‒cell communication (Jin *et al*, 2021). The ligand‒receptor interaction database was included in the package.

### Protein‒Protein Interaction Analysis

To determine the core DEGs at each age, we used the online tool STRING (version 12.0) (http://string-db.org/) to construct PPI networks, and the parameters were set to the default value.

### Cell culture and the generation of SBMA OL model cells

Tissue culture media and reagents were purchased from Invitrogen. Oli-neu cells were kindly provided by Prof. J. Trotter (University of Mainz) and cultured in Sato medium containing Dulbecco’s modified Eagle’s medium (DMEM) supplemented with 1% (v/v) horse serum, insulin (5 μg/ml), penicillin (100 U/ml), streptomycin (0.1 mg/ml), 1% (v/v) N2 supplement, sodium selenite (190 nM), gentamicin (25 μg/ml), triiodothyronine (400 nM) and L-thyroxine (520 nM). Cells were kept in a humidified atmosphere with 5% CO2 at 37°C. Lentiviruses expressing full-length human AR containing 17 or 97 CAG repeats were introduced into Oli-neu cells, thereby generating cell lines harboring AR-17Q as control cells and cell lines harboring AR-97Q as SBMA model cells. Differentiation was induced by adding 1 μM PD174265 (Calbiochem, CA, USA) to the medium for 72 hours (Naffaa *et al*, 2022).

### Lentiviral production

Lentivirus was prepared following Campeu’s protocols (Campeau *et al*, 2009). Briefly, lentiviral particles were produced in HEK293T cells by transfection using Lipofectamine 2000 (#11668030, Thermo Fisher Scientific). Lentivirus-containing supernatant was collected at 72 hours after transfection and stored at −80°C. The viral titer was measured using a Lenti-X qRT‒PCR Titration Kit (#631235, TaKaRa, Shiga, Japan).

### Bulk RNA-seq

Total RNA was extracted from the SBMA OL cell model and the control cell model using the RNeasy Mini Kit (Qiagen, Hilden, Germany). The quality and quantity of the RNA obtained were checked using a NanoDrop 2000 spectrophotometer (Thermo Fisher Scientific) and analyzed with the Agilent 2100 Bioanalyzer system (Agilent Technologies Inc.). The RNA was sent to Macrogen (Macrogen Inc., Seoul, South Korea) for library preparation and sequencing. The sequencing of the 8 libraries was carried out using the NovaSeq sequencing protocol and TruSeq stranded mRNA Library Kit, following a paired-end 100 bp strategy on the Illumina NovaSeq 6000 platform.

### Quantitative RT‒PCR

Total RNA was extracted from the cells using the RNeasy Mini Kit (Qiagen). The extracted RNA was then reverse-transcribed into first-strand cDNA using the ReverTra Ace qPCR RT Kit (Toyobo, Osaka, Japan). RT‒PCR was performed using a KOD SYBR qPCR kit (Toyobo), and the amplified products were detected with a CFX Connect system (Bio-Rad Laboratories, Hercules, CA, USA). The reaction conditions were as follows: 98.0 °C for 2 min, 45 cycles of 10 s at 98.0 °C, 10 s at 60.0 °C and 30 s at 68.0 °C. The expression level of the internal control, β_2_ microglobulin, was simultaneously quantified. The primers are listed in Supplementary Table 1.

### Immunoblotting

Cultured cells were lysed in buffer containing 50 mM Tris-HCl (pH 8.0), 150 mM NaCl, 1% Nonidet P-40, 0.5% deoxycholate, 0.1% SDS, and 1 mM 2-mercaptoethanol with the Halt Protease and Phosphatase Inhibitor Cocktail (Thermo Fisher Scientific). We separated equal amounts of protein on 5-20% SDS‒PAGE gels (Wako, Osaka, Japan) and transferred them to Hybond-P membranes (GE Healthcare, Piscataway, NJ, USA). The following primary antibodies and dilutions were used: AR (#5153, 1:2000; Cell Signaling Technology), Mog (ab233549, 1:2000; Abcam), PDGFRα (#3164S, 1:2000; Cell Signaling Technology), and Sox10 (sc-365692, 1:200; Santa Cruz). Primary antibodies bound to the proteins were probed with a 1:5000 dilution of horseradish peroxidase-conjugated secondary antibodies, and the bands were detected using an immunoreaction enhancing solution (Can Get Signal; Toyobo) and enhanced chemiluminescence (ECL Prime; GE Healthcare). Chemiluminescence signals were digitized using a ChemiDoc MP imaging system (Bio-Rad Laboratories). Membranes were reprobed with an anti-GAPDH antibody (ab8245, 1:5000; Abcam) for normalization.

### Immunohistochemistry

Mouse tissues were dissected, postfixed in 10% phosphate buffered formalin, and paraffin-embedded. Six-micron-thick sections were prepared from the paraffin-embedded tissues. The sections designated to be stained with the anti-polyglutamine antibody (1C2) (MAB1574, 1:20,000; Millipore) were treated with formic acid for 5 min at room temperature. The sections designated to be incubated with the human AR antibody (#5153, 1:1000, Cell Signaling Technology) were boiled in 10 mM citrate buffer for 15 min. Primary antibodies bound to proteins were incubated with a secondary antibody labeled with a polymer as part of the Envision + system containing horseradish peroxidase (Dako Cytomation, Gostrup, Denmark). Images of immunohistochemically stained sections were obtained using an optical microscope (BX51, Olympus, Tokyo, Japan). Immunoreactivity and cell size were analyzed using ImageJ software (NIH, Bethesda, MD). The mean ± standard error of the mean (SEM) of the obtained values is presented in arbitrary units.

### Immunofluorescence for mouse tissues and human autopsy samples

Mouse tissues and autopsied human spinal cords were dissected, postfixed with 10% phosphate-buffered formalin and processed for paraffin embedding. Six-micron-thick sections were prepared from paraffin-embedded tissues. The sections designated to be stained with the anti-polyglutamine antibody (1C2) (MAB1574, 1:20,000; Millipore) were treated with formic acid for 5 min at room temperature. The sections designated to be incubated with the anti-human AR (554224, 1:100; Biosciences), Plp1 (ab28486, 1:500; Abcam), Olig2 (#18953, 1:100; IBL), Sox10 (MAB2864, 1:50; R&D for human samples, sc-365692, 1:200; Santacruz for mouse samples), Mbp (ab7349, 1:200; Abcam), Asic2 (ASC-012, 1:100; alomone labs), and Fam155a (orb2222, 1:100; biorbyt) antibodies were boiled in 10 mM citrate buffer for 15 min. After washing, the samples were incubated with Alexa-488–conjugated donkey anti-mouse IgG (1:1000; A21202, Invitrogen), Alexa-546– conjugated goat anti-mouse IgG (1:1000; A11003, Invitrogen), Alexa-488–conjugated goat anti-rabbit IgG (1:1000; A11008, Invitrogen), or Alexa-568–conjugated goat anti-rabbit IgG (1:1000; A11036, Invitrogen) for 1 hour, stained with Hoechst 33342 (H3570, Invitrogen), mounted using ProLong Diamond Antifade Mountant (P36961, Invitrogen), then imaged with a fluorescence microscope (BZ-X810, Keyence Corporation, Osaka, Japan).

### Statistical analysis

GraphPad Prism 9.0 software was used for statistical analysis and the plotting of statistical graphs. Data are expressed as the mean ± SEM. Differences between the two groups were evaluated by a two-tailed Student’s t test. The results were considered statistically significant at p < 0.05. Principal component analysis (PCA) and heatmap generation, as well as enriched pathway analysis of microarray data were performed using iDEP.94(Ge *et al*, 2018). Upregulated DEGs in the OLs of AR-97Q mice (log_2_FC > 0.1) at 6 weeks and the upregulated genes in AR-97Q cells compared to AR-24Q cells were analyzed with the Metascape portal (www.Metascape.org) (Zhou *et al*, 2019).

### Study approval

All animal experiments were performed in accordance with the National Institutes of Health Guide for the Care and Use of Laboratory Animals and with the approval of the Nagoya University Animal Experiment Committee. The collection of autopsied human tissues and their use in this study were approved by the Ethics Review Committee of Nagoya University Graduate School of Medicine. Experimental procedures involving human subjects were conducted in accordance with the Declaration of Helsinki, the Ethical Guidelines for Medical and Biological Research Involving Human Subjects endorsed by the Japanese government.

## Data availability

Single-nucleus RNA sequencing data in this paper is available in the NCBI Gene Expression Omnibus (GEO) database with accession number GSE248684 [https://www.ncbi.nlm.nih.gov/geo/query/acc.cgi?acc=GSE248684]. Bulk RNA sequencing data in this paper is available in the NCBI GEO database with accession number GSE247374 [https://www.ncbi.nlm.nih.gov/geo/query/acc.cgi?acc=GSE247374]. Other data in this study are provided in the Source Data file.

## Code availability

The codes used for data analyses in this study have been deposited on GitHub under the following link: [https://github.com/madoka-iida/Iida-et-al.-2023].

## Author contributions

Project planning was performed by M.I., K.S., and M.K.; experiments were performed by M.I., K.S., T.H., K.S., K.M., T.A, and M.K.; and data were analyzed by M.I., K.S., Y.O., M.I., K.H., and M.K. The first draft of the manuscript was prepared by M.I. and M.K.; the manuscript layout was designed by M.K.

## Acknowledgments

We acknowledge the Japan Research Activity Support, Inc. (JRAS Inc.) for setting up the pipelines for scRNA-Seq analysis. This work was supported by JSPS KAKENHI Grant Numbers JP21K20686 and JP22K15705 to MI, JP20H00527 and JP23H00420 to MK; JP22nk0101575 to KS, and JP22K15706 to TH; AMED under Grant Numbers JP23bm1423003 to MK and JP22am0401007 to KS; a grant from Takeda Science Foundation (MI), and a grant from the Japanese SBMA Patient Group Research Support Program (MI). No other agencies provided funding, and the investigators had sole discretion over the study design; collection, analysis, and interpretation of data; writing of the report; and the decision to submit it for publication.

---

**A table and it’s legend**

**Supplementary Table 1.**
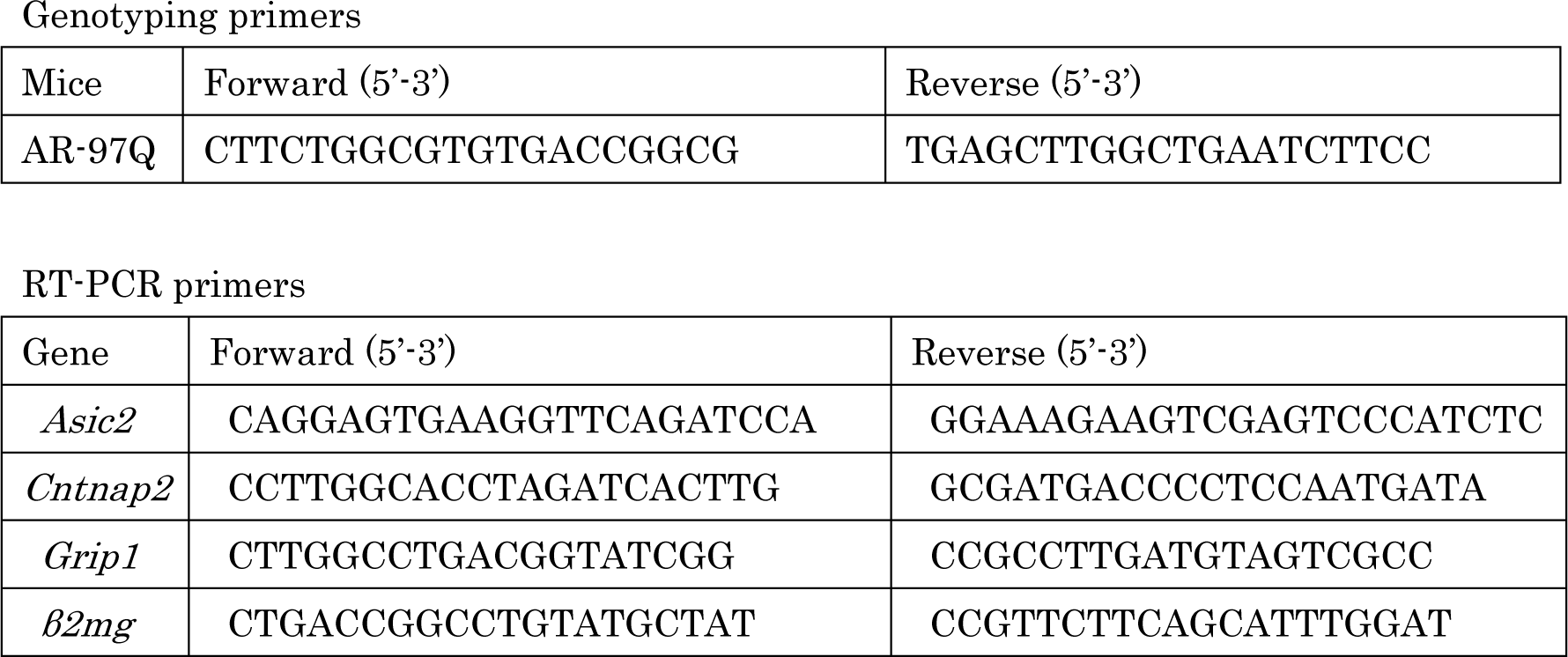
List of used primers.

## Expanded View Figure legends

**Supplementary Fig. 1. Summary of snRNA-seq**

**A,** Number of genes, number of UMI sequences, and proportion of UMI sequences from mitochondrial genes (from left to right) per cell in each sample. **B,** The average number of nuclei of each sample before quality control filtering. **C,** Number of cells for each sample after quality control filtering.

**Supplementary Fig. 2. snRNA-seq distinguishes major cell types in spinal cord**

**A,** The dot plots of known cell marker genes in each cell type. **B,** The dot plots of known cell marker genes related to oligodendrocyte (OL) differentiation in OL, oligodendrocyte precursor cells (OPC), and OL progenitors.

**Supplementary Fig.3. Impaired oligodendrocytes in spinal cords of AR-97Q mice** Immunohistochemical analysis of myelin basic protein (Mbp) in the spinal cords of wild-type and AR-97Q mice at 13 weeks. Scale bar: 50μm.

**Supplementary Fig. 4. Microarray analysis of spinal cords at early stage of disease**

**A,** The results of principal component analysis (PCA) for each sample. **B,** Number of differentially expressed genes (FDR < 0.1, FC > 1.5). Red indicates upregulated genes in AR-97Q mice and blue indicates downregulated genes in AR-97Q mice compared to AR-24Q mice. **C,** Heatmap showing results of hierarchical clustering analysis of genes that are significantly altered. **D,** Enriched pathways in DEGs for AR-97Q mice. **E,** Relative expression levels of channel related genes: *Asic2*, *Fam155a*, and *Kcnip4* in each mouse. **F,** Relative expression levels of genes related to synaptic function: *Cntnap2*, *Nrxn3*, and *Grip1* in each mouse.

**Supplementary Fig. 5. Unique gene expression signatures of oligodendrocyte cluster at 9 weeks**

**A,** t-Distributed stochastic neighbor embedding (t-SNE) and uniform manifold approximation and projection (UMAP) plot visualizing clusters of single nuclei in the spinal cord of AR-97Q and wild-type mice at 9 weeks. **B,** t-SNE plot of the oligodendrocyte (OL) cluster colored with the genotype: orange dots represent AR-97Q mice (A9) and green dots represent wild-type mice (W9). **C,** Volcano plot showing DEGs of OL cluster of AR-97Q mice and wild-type mice. The top 5 genes and last 5 genes are marked. **D,** Predicted protein interaction (PPI) networks for the top 20 upregulated genes in AR-97Q mice. Genes colored in red have cation channel complex and genes colored in purple have synapse in the cellular component of GO terms. **E,** PPI networks for the top 20 downregulated genes in AR-97Q mice. Genes colored in red are related to myelin sheath. **F, G,** The enrichment of the top 100 upregulated genes in AR-97Q mice in the biological process (**F**) and molecular function (**G**) categories (log_2_FC > 0.338). **H, I,** The enrichment of the top 100 downregulated genes in AR-97Q mice in the biological process (**H**) and molecular function (**I**) categories (log_2_FC < −0.234). A9, AR-97Q mice at 9 weeks; W9, wild-type mice at 9 weeks.

**Supplementary Fig. 6. Unique gene expression signatures of OL cluster at 3 weeks**

**A,** t-Distributed stochastic neighbor embedding (t-SNE) and uniform manifold approximation and projection (UMAP) plot visualizing clusters of single nuclei in the spinal cord of AR-97Q and wild-type mice at 3 weeks. **B,** t-SNE plot of the oligodendrocyte (OL) cluster colored with the genotype: orange dots represent AR-97Q mice (A3) and green dots represent wild-type mice (W3). **C,** Volcano plot showing DEGs of oligodendrocyte (OL) cluster of AR-97Q mice and wild-type mice. The top 5 genes and last 5 genes are marked. **D,** Predicted protein interaction (PPI) networks for the top 17 upregulated genes in AR-97Q mice. Genes colored in red have cation channel complex and genes colored in purple have synaptic membrane in cellular component of GO terms. **E,** PPI networks for the top 20 downregulated genes in AR-97Q mice. Genes colored in red are related to axons. **F, G,** The GO enrichment of 18 upregulated genes in AR-97Q mice in the biological process (**F**) and molecular function (**G**) categories (log_2_FC > 0.22). **H, I,** The enrichment of 67 downregulated genes in AR-97Q mice in the biological process (**H**) and molecular function (**I**) categories (log_2_FC < −0.2). A3, AR-97Q mice at 3 weeks; W3, wild-type mice at 3 weeks.

**Supplementary Fig. 7. Oligodendrocyte heterogeneity at 3 weeks**

**A,** Uniform manifold approximation and projection (UMAP) plot of the oligodendrocyte (OL) cluster colored with the genotype: orange dots represent AR-97Q mice (A3) and green dots represent wild-type mice (W3). **B,** UMAP plot of the OL cluster of 3 weeks of age with the associated cell subcluster. **C,** Proportion of each cell subcluster in AR-97Q and wild-type mice. **D,** The enrichment of the Reactome pathway of each subcluster. Red arrows indicate subcluster 0, a cluster dominant in AR-97Q mice.

**Supplementary Fig. 8. PPI networks of transcription factors**

Predicted protein interaction (PPI) networks of transcription factors that are related to top 10 and last 10 DEGs of oligodendrocytes of AR-97Q mice at 3 weeks compared to those of wild-type mice at 3 weeks.

**Supplementary Fig. 9. Pseudotime analysis based on clusters of OL lineage cells of wild-type mice**

**A,** Pseudotime analysis inferred from clusters of oligodendrocyte (OL) lineage cells of wild-type mice at 3, 6, 9, and 13 weeks. **B,** Uniform manifold approximation and projection (UMAP) visualization of clusters of OL lineage cells colored by weeks of age. **C,** Boxplot showing the distribution of pseudotime within each sample. Vertical bars indicate median values.

**Supplementary Fig. 10. Pseudotime kinetics of *Asic2* and *Fam155a* in oligodendrocyte lineage cells of AR-97Q and wild-type mice**

**A,** Uniform manifold approximation and projection (UMAP) visualization of clusters of oligodendrocyte (OL) lineage cells of wild-type mice at 4 stages colored by Seurat package (resolution = 1.2). **B, C,** Pseudotime kinetics of *Asic2* (**B**), and *Fam155a* (**C**).

**Supplementary Fig. 11. Expression of Asic2 in oligodendrocytes of spinal cords of AR-97Q mice**

**A, B,** Immunohistochemical analysis of of spinal cords of wild-type mice and AR-97Q mice at 6 weeks (**A**) and 13 weeks (**B**). Scale bars: 50 μm.

**Supplementary Fig. 12. Expression of Fam155a in oligodendrocytes of spinal cords of AR-97Q mice**

**A, B,** Immunohistochemical analysis of spinal cords of wild-type mice and AR-97Q mice at 6 weeks (**A**) and 13 weeks (**B**). Scale bars: 50 μm.

**Supplementary Fig. 13. Top DEGs that are common across cell types**

**A,** Common DEGs upregulated in AR-97Q mice in oligodendrocytes (OLs) and astrocytes (left), and in OLs, inhibitory neurons, and excitatory neurons (right) at 3 weeks. **B,** Common top 20 DEGs across OLs, astrocytes, and microglia at 6 weeks. Numbers in parentheses indicate the number of DEGs.

**Supplementary Fig. 14. Disease-dependent DEGs of OPC at 6 and 13 weeks**

**A,** Volcano plot showing differential gene expression of the oligodendrocyte precursor cells (OPC) cluster of AR-97Q mice and wild-type mice at 6 weeks. The top 5 genes and last 5 genes are marked. **B,** The enrichment of 18 upregulated genes in AR-97Q mice at 6 weeks in the biological process category (log_2_FC > 0.37). **C,** Volcano plot showing differential gene expression of OPC cluster of AR-97Q mice and wild-type mice at 13 weeks. The top 5 genes and last 5 genes are marked. **D,** The enrichment of 100 downregulated genes in AR-97Q mice at 13 weeks in the biological process category (log_2_FC < −0.368).

**Supplementary Fig. 15. Comparison of the RNA-seq data of oligodendrocyte cell model of SBMA and OL data from snRNA-seq**

**A,** Ratio of the number of common genes upregulated or downregulated in both cell and mouse model versus the number of total genes that were measured the expression levels by both RNA-seqs. **B,** GO terms (-log_10_(P) > 5) of genes that are among DEGs upregulated in OL of AR-97Q mice (log_2_FC > 0.1) at 6 weeks and among genes upregulated in AR-97Q cells compared to AR-24Q cells. \**p* < 0.001, equality of proportions hypothesis test.

